# RS-fMRI Evidence for Differential Within- and Between-Module Neural Interactions Across Age

**DOI:** 10.1101/2025.07.08.663619

**Authors:** Tien-Wen Lee

**Author notes:** Corresponding author at The NCI Clinical Research Foundation, New Jersey, USA Address: 111 Howard Blvd., Suite 204, Mt. Arlington, NJ 07856, Web: http://neuroci.com.

## Abstract

**Background:** Previous resting-state functional MRI (rsfMRI) studies have identified a robust inverse relationship between nodal strength and amplitude of low-frequency fluctuations (ALFF) across cortical modules. This study examined whether this negative relationship exists within modules and further explored age-related effects on these associations.

**Methods:** Using MOSI (modularity analysis and similarity measurements), rsfMRI data from three public datasets spanning different age cohorts were analyzed. Correlations between ALFF and nodal strength between modules or voxel concordance within modules were calculated. Geometric mean p-values assessed robustness at the individual level.

**Results:** Confirming prior findings, a significant inverse correlation between nodal strength and ALFF was observed at the between-module level (geometric p-values 10^−4^ to 10^−5^). Within-module negative associations were nonsignificant in younger cohorts at individual level (mean ages 10 and 21) but became significant in the older cohort (mean age 33). The magnitude of this negative association increased with age, consistent with maturation of local inhibitory network mechanisms.

**Conclusions:** The findings support MOSI as a valid FP method for cortical network construction, with within-module inhibitory effects strengthening in adulthood. The age-dependent modulation reveals network maturation at the modular level, with implications for neurodevelopmental and neuropsychiatric conditions.

## Introduction

Network analysis has emerged as a central theme in brain science research. Neuroimaging studies suggest that local neural activity may be influenced by the network environment in which a neural node is embedded (Lee and Xue, 2017; Misic et al., 2011). When using resting-state functional magnetic resonance imaging (rsfMRI) to investigate network properties, a major challenge arises from the sheer number of voxels (typically tens to hundreds of thousands), which can render computation intractable. To address this, cortical parcellation based on anatomical atlases has been widely applied in fMRI research to reduce data dimensionality to a more manageable set of regions. Nevertheless, some researchers argue for voxel-wise network analysis to preserve fine-grained information (Stanley et al., 2013).

Recently, functional parcellation (FP) methods for rsfMRI have been developed that cluster voxels with similar temporal dynamics into neural nodes, relying on data-driven rather than atlas-based definitions (Shen et al., 2010; Wig et al., 2014). The author introduced MOSI (modularity analysis and similarity measurements), a novel FP approach applied in several psychiatric studies (Lee, 2025a, b, c; Lee and Tramontano, 2021). Using MOSI, a robust inverse relationship between nodal strength and the amplitude of low-frequency fluctuations (ALFF) has been identified in multiple public datasets (Lee, 2025a, b, c; Lee and Tramontano, 2021, 2025), suggesting that cortical networks may exert a net inhibitory influence on individual neural nodes.

This study aimed to test whether this inverse relationship, previously observed across modules, also extends to interactions within modules. Specifically, it examined whether voxel concordance within a module—where higher concordance indicates stronger “nodal strength” from intra-module voxels—exhibits a similar inhibitory association with average ALFF. Failure to replicate this relationship would empirically support FP as a valid preparatory step for network construction, given the distinct physiological characteristics of local versus global interactions. Finally, age-related effects on these network properties were explored from three public datasets spanning adolescence to adulthood.

## Methods

### Subjects, MRI data, preprocessing and functional parcellation

MRI data were obtained from healthy control groups (n = 60 each) of three youth and adult datasets: ABIDE (mean age 21.0 ± 4.2 years) (Di Martino et al., 2014), ADHD-200 (mean age 10.1 ± 1.7 years) (Bellec et al., 2017), and CAN-BIND (BrainCode) (mean age 32.7 ± 11.6 years) (Lam et al., 2016). Functional and structural MRI were acquired using 3.0 Tesla scanners with whole-brain coverage and comparable parameters (TR = 1.5–2 s, voxel size ≈ 3×3×4 to 4×4×4 mm^3^, volumes 170–300). Detailed acquisition protocols are provided in the original references.

MRI data from the healthy control groups of the three datasets were directly obtained from the author’s previous analyses (Lee, 2025a, b, c). Preprocessing and functional parcellation followed the established pipelines: (1) EPI data were processed using AFNI (Cox, 1996), including despiking, slice-timing correction, motion realignment, T1 anatomical registration, spatial smoothing, and bandpass filtering (0.01–0.10 Hz). Mean ALFF was computed for each MOSI-defined module or partition, reflecting the average low-frequency power of the BOLD signal within these regions; (2) Functional parcellation employed MOSI, integrating modular analysis and similarity measures through iterative split-merge steps based on spatial proximity and functional similarity. The Louvain algorithm was used with Gamma parameters of 0.65, 0.75, and 0.85 to generate parcellations at multiple resolutions (Lee and Tramontano, 2021).

### Nodal Strength–ALFF Relationship in the Between-Module Condition

The analysis of regional and inter-regional interactions was motivated by the observation that a node’s global connectivity—quantified as nodal strength in weighted graphs (i.e., the sum or average of connection weights centered on a node)—can modulate its local neural activity (Lee and Tramontano, 2025; Lee and Xue, 2017). In particular, nodal strength at lower frequency ranges of neural signals (e.g., fMRI and delta/theta bands in EEG) has been linked to inhibitory network influences (Lee and Tramontano, 2025). To correct for variations in module size, a weighting strategy was implemented based on the square root of voxel count, as described in previous work (Lee, 2025b).

### Local Concordance and ALFF in the Within-Module Condition

The relationship between within-module concordance and ALFF has not been systematically explored. The author examined the association between ALFF and Kendall’s coefficient of concordance (Kendall’s W) within functionally defined cortical modules (at specific Gamma values), using partial correlation to control for module size. This adjustment was necessary because smaller modules inherently tend to exhibit higher Kendall’s W, potentially introducing bias. In addition, mean correlation coefficients across all voxel pairs within each module (N×(N−1)/2) were computed as an alternative measure of concordance. Since these results closely mirrored those obtained with Kendall’s W, they are not reported separately.

### Statistical comparisons

The correlation coefficients were Fisher z-transformed, and one-sample t-tests were performed for the between-module and within-module analyses. To evaluate the robustness of associations at the individual level, a representative P value—termed the *geometric P*—was computed as the geometric mean of individual subject P values. For each functional partition (specific Gamma value), ALFF and nodal strength correlations across modules yielded 60 P values per dataset (p_1_, p_2_,, p_60_) ^1/ 60^, with the geometric P calculated as (p_1_ × p_2_ × … × p _60_)^1/60^. The same procedure was applied to within-module concordance–ALFF correlations. Notably, while t-test P values may achieve statistical significance under consistently negative correlations, such group-level effects do not necessarily reflect robust associations at the individual level. Given two conditions across three Gamma values (six comparisons per dataset), and the partial dependence of Gamma outputs, a threshold of p<0.01 (0.05/5) was adopted to account for multiple comparisons.

## Results

The robust inverse relationship between neural metrics and ALFF at the individual level was reflected in geometric P values ranging from 10^−4^ to 10^−5^, as reported in previous studies (Lee, 2025a, b, c; Lee and Tramontano, 2025). Similar to the between-module findings, a negative association was observed between within-module concordance and ALFF, becoming markedly stronger with age. The corresponding geometric P values for the within-module condition were not significant in ADHD-200 and ABIDE (younger cohorts; mean ages 10 and 21 years, respectively), but reached significance in CAN-BIND (older cohort; mean age 33 years). These results are summarized in Table 1.

**Table 1.** Between-and Within-Module Associations Between ALFF and Network Measures

| Gamma | Between-Module |  |  |  |  | With-Module |  |  |  |  |
| --- | --- | --- | --- | --- | --- | --- | --- | --- | --- | --- |
|  | mean | std | t-score | pval | geo_pval | mean | std | t-score | pval | geo_pval |
| ABIDE |  |  |  |  |  |  |  |  |  |  |
| 0.65 | -0.29 | 0.15 | -13.88 | 3.05E-20 | 1.35E-04* | -0.06 | 0.15 | -3.06 | 0.003 | 0.169 |
| 0.75 | -0.25 | 0.12 | -15.13 | 5.65E-22 | 1.84E-04* | -0.15 | 0.14 | -8.05 | 4.42E-11 | 0.050 |
| 0.85 | -0.21 | 0.10 | -15.45 | 2.12E-22 | 4.28E-04* | -0.15 | 0.14 | -7.75 | 1.43E-10 | 0.035 |
| ADHD-200 |  |  |  |  |  |  |  |  |  |  |
| 0.65 | -0.25 | 0.20 | -9.96 | 2.99E-14 | 8.83E-04* | -0.02 | 0.15 | -1.29 | 0.203738 | 0.271 |
| 0.75 | -0.23 | 0.16 | -11.40 | 1.49E-16 | 5.44E-04* | -0.08 | 0.15 | -3.97 | 2.00E-04 | 0.119 |
| 0.85 | -0.19 | 0.14 | -10.27 | 9.52E-15 | 8.32E-04* | -0.11 | 0.15 | -5.69 | 4.26E-07 | 0.069 |
| CAN-BIND |  |  |  |  |  |  |  |  |  |  |
| 0.65 | -0.43 | 0.18 | -15.87 | 5.94E-23 | 1.98E-05* | -0.37 | 0.19 | -14.29 | 8.04E-21 | 0.003* |
| 0.75 | -0.39 | 0.17 | -15.88 | 5.75E-23 | 1.45E-05* | -0.39 | 0.19 | -15.41 | 2.36E-22 | 0.0008* |
| 0.85 | -0.34 | 0.15 | -15.82 | 6.93E-23 | 4.07E-05* | -0.41 | 0.16 | -18.74 | 1.59E-26 | 0.0002* |

Between-module associations were computed as the correlation between nodal strength and ALFF, while within-module associations were computed as the correlation between Kendall's coefficient of concordance (Kendall's W) and ALFF. The table reports mean correlation (Fisher z-transformed), standard deviation (std), t-scores (t-score), uncorrected p-values (pval), and geometric p-values (geo\_pval) for each of the three datasets (ABIDE, ADHD200, CAN-BIND) across different Gamma values. Asterisks (\*) indicate geo\_pval < 0.01.

## Discussion

This report investigated whether the inverse association between nodal strength and ALFF observed in the between-module context (left panel of Table 1) also holds within modules (right panel of Table 1). While a negative association was replicated, the geometric P values were nonsignificant in the younger cohorts. In contrast, a significant negative correlation emerged in the older cohort, though with lower statistical robustness compared to the between-module scenario. These findings support FP as a valid preparatory step for network construction and subsequent analyses. They also partly align with Stanley et al.’s assertion that a voxel can function as an independent neural node; a property that seems to develop only after early adulthood beyond the twenties (Stanley et al., 2013).

The cortex—especially the prefrontal regions—undergoes prolonged structural and functional maturation during adolescence, characterized by incomplete network differentiation and integration (Konrad et al., 2013). In adulthood, increased network integration and efficiency become established as cortical circuits stabilize, reflecting reduced plasticity alongside enhanced modular specialization. During adolescence, excitatory glutamatergic neurotransmission predominates, while inhibitory GABAergic systems remain immature, resulting in heightened neural excitability and less regulated cortical activity (Arain et al., 2013). With maturation into adulthood, strengthened GABAergic inhibitory circuits promote a more balanced excitation–inhibition ratio, supporting improved cortical stability, functional differentiation, and refined local network interactions.

This developmental trajectory summarized above aligns with the present finding of a significant negative correlation between within-module voxel concordance (an approximation of nodal strength) and ALFF in adults. Stronger local synchronization appears linked to reduced neural activity, consistent with increasing inhibitory influences within local networks during maturation. Furthermore, the emergence of a significant within-module inhibitory effect in older individuals indicates that voxels within a module begin to demonstrate functional autonomy, behaving more like independent neural nodes relative to their surrounding network—paralleling the between-module organization (Stanley et al., 2013). This enhanced within-module inhibition may also contribute to the reduced variability of neural activity seen with aging (Garrett et al., 2011).

Future studies incorporating elderly populations, beyond the age groups examined here (10s, 20s, and 30s), as well as gender-specific analyses, will require larger samples to clarify these effects. Moreover, the differentiation between within- and between-module manifestations may hold promise as a novel neural marker for investigating the neuropathology of diverse neuropsychiatric conditions.

## Authors Contributions

This report is authored by a single individual.

## Acknowledgments

The author would like to thank the Autism Brain Imaging Data Exchange (ABIDE), coordinated by New York University Langone Medical Center and supported by the Child Mind Institute, for providing access to MRI datasets collected through the collaborative efforts of multiple international sites. Special thanks also to the ADHD-200 Consortium, organized by the Child Mind Institute, New York University Langone Medical Center, Kennedy Krieger Institute, Oregon Health & Science University, and other partners for generously sharing MRI data from diverse sites worldwide. In addition, the author gratefully acknowledge the Canadian Biomarker Integration Network for Depression (CAN-BIND), the Ontario Brain Institute, the Brain-CODE platform, and the Government of Ontario, as well as all collaborators listed at [https://www.canbind.ca/about-can-bind/our-team/executive-committee/], for their contributions to data sharing and coordination.

## Statements and Declarations

No conflicts of interest to declare.

## Compliance with ethical standards

This research analyzed the data from publicly released datasets. The author carried out no animal or human studies for this article.

## References

Arain, M., Haque, M., Johal, L., Mathur, P., Nel, W., Rais, A., Sandhu, R., Sharma, S. (2013) Maturation of the adolescent brain. Neuropsychiatr Dis Treat 9, 449–461.

Bellec, P., Chu, C., Chouinard-Decorte, F., Benhajali, Y., Margulies, D.S., Craddock, R.C. (2017) The Neuro Bureau ADHD-200 Preprocessed repository. Neuroimage 144, 275–286.

Cox, R.W. (1996) AFNI: software for analysis and visualization of functional magnetic resonance neuroimages. Comput Biomed Res 29, 162–173.

Di Martino, A., Yan, C.G., Li, Q., Denio, E., Castellanos, F.X., Alaerts, K., Anderson, J.S., Assaf, M., Bookheimer, S.Y., Dapretto, M., Deen, B., Delmonte, S., Dinstein, I., Ertl-Wagner, B., Fair, D.A., Gallagher, L., Kennedy, D.P., Keown, C.L., Keysers, C., Lainhart, J.E., Lord, C., Luna, B., Menon, V., Minshew, N.J., Monk, C.S., Mueller, S., Muller, R.A., Nebel, M.B., Nigg, J.T., O’Hearn, K., Pelphrey, K.A., Peltier, S.J., Rudie, J.D., Sunaert, S., Thioux, M., Tyszka, J.M., Uddin, L.Q., Verhoeven, J.S., Wenderoth, N., Wiggins, J.L., Mostofsky, S.H., Milham, M.P. (2014) The autism brain imaging data exchange: towards a large-scale evaluation of the intrinsic brain architecture in autism. Mol Psychiatry 19, 659–667.

Garrett, D.D., Kovacevic, N., McIntosh, A.R., Grady, C.L. (2011) The importance of being variable. J Neurosci 31, 4496–4503.

Konrad, K., Firk, C., Uhlhaas, P.J. (2013) Brain development during adolescence: neuroscientific insights into this developmental period. Dtsch Arztebl Int 110, 425–431.

Lam, R.W., Milev, R., Rotzinger, S., Andreazza, A.C., Blier, P., Brenner, C., Daskalakis, Z.J., Dharsee, M., Downar, J., Evans, K.R. (2016) Discovering biomarkers for antidepressant response: protocol from the Canadian biomarker integration network in depression (CAN-BIND) and clinical characteristics of the first patient cohort. BMC Psychiatry 16, 1–13.

Lee, T.W. (2025a) Asymmetrical frontal–subcortical aberrance in ADHD: empirical support of a developmental network model medRxiv.

Lee, T.W. (2025b) Framing major depressive disorder as a condition of network imbalance at the compartment level: A proof-of-concept study. Cerebral Cortex 35, bhaf089.

Lee, T.W. (2025c) RS-fMRI evidence of left frontal lobe developmental deviation as a potential core pathology of autism spectrum disorder. In Review.

Lee, T.W., Tramontano, G. (2021) Automatic parcellation of resting-state cortical dynamics by iterative community detection and similarity measurements. AIMS Neurosci 8, 526–542.

Lee, T.W., Tramontano, G. (2025) Inverse relationship between nodal strength and nodal power: Insights from separate resting fMRI and EEG datasets. Journal of Neuroscience Methods (accepted) 418, 110438

Lee, T.W., Xue, S.W. (2017) Linking graph features of anatomical architecture to regional brain activity: A multi-modal MRI study. Neuroscience Letters 651, 123–127.

Misic, B., Vakorin, V.A., Paus, T., McIntosh, A.R. (2011) Functional embedding predicts the variability of neural activity. Front Syst Neurosci 5, 90.

Shen, X., Papademetris, X., Constable, R.T. (2010) Graph-theory based parcellation of functional subunits in the brain from resting-state fMRI data. Neuroimage 50, 1027–1035.

Stanley, M.L., Moussa, M.N., Paolini, B.M., Lyday, R.G., Burdette, J.H., Laurienti, P.J. (2013) Defining nodes in complex brain networks. Front Comput Neurosci 7, 169.

Wig, G.S., Laumann, T.O., Cohen, A.L., Power, J.D., Nelson, S.M., Glasser, M.F., Miezin, F.M., Snyder, A.Z., Schlaggar, B.L., Petersen, S.E. (2014) Parcellating an individual subject’s cortical and subcortical brain structures using snowball sampling of resting-state correlations. Cereb Cortex 24, 2036–2054.

